# An fMRI-based brain marker predicts individual differences in delay discounting

**DOI:** 10.1101/2021.03.18.435969

**Authors:** Leonie Koban, Sangil Lee, Daniela S. Schelski, Marie-Christine Simon, Caryn Lerman, Bernd Weber, Joseph W. Kable, Hilke Plassmann

**Affiliations:** Marketing Area, INSEAD, Fontainebleau, France; Control-Interoception-Control Team, Paris Brain Institute (ICM), INSERM U 1127, CNRS UMR 7225, Sorbonne University, Paris, France; Department of Psychology, University of Pennsylvania, Philadelphia, PA, USA; Center for Economics and Neuroscience, University of Bonn, Bonn, Germany; Institute of Experimental Epileptology and Cognition Research, University of Bonn, Germany; Institute for Nutrition Science, University of Bonn, Bonn, Germany; Norris Comprehensive Cancer Center, University of Southern California, Los Angeles, CA, USA

## Abstract

Individual differences in impatience—how much we discount future compared to immediate rewards—are associated with general life outcomes and related to substance use, psychiatric diseases, and obesity. Here, we use machine-learning on fMRI activity during an intertemporal choice task to develop a brain marker of individual differences in delay discounting. Study 1 (N=110) was used as a training and cross-validation set, resulting in significant prediction accuracy (*r* = 0.49) and suggesting an interplay between brain regions associated with affect, value, and cognitive control. The validity of the brain marker was replicated in an independent data set (Study 2, N=145, *r* = 0.45). In both studies, responses of the marker significantly differed between overweight and lean individuals. This pattern is a first step towards a generalizable neuromarker of delay discounting and a potentially transdiagnostic phenotype, which can be used as a brain-based target measure in future studies.

## Introduction

Many decisions in daily life require choices that have consequences at different points in time. For example, most people need to decide at the end of the month whether to put part of their paycheck towards a retirement fund or directly spend it on something fun like a short vacation. Those trade-offs between choosing an option that is immediately rewarding versus one that will be more rewarding in the long run are hard and people differ substantially in their ability to delay immediate gratification. Both intuition and accumulating evidence suggests that *patience for rewards*—the ability to make choices that are aligned with long-time goals and to resist immediate gratification—is closely related to other advantageous behaviors, better health, and overall life achievement (Mischel et al., 1988; Moffitt et al., 2011).

In economics, patience for rewards is typically measured as *delay discounting* (DD), which quantifies how much people discount future compared to immediate rewards (Frederick et al., 2002; Kirby and Herrnstein, 1995; Laibson, 1997). Most people prefer sooner rather than later rewards (e.g., almost everybody would prefer 100 Euro right now over 101 Euro in 2 years). Yet, the degree to which delayed rewards are discounted varies widely between individuals. Greater delay discounting (i.e., greater impatience) is associated with smoking, obesity and overweight (Amlung et al., 2016; Bickel et al., 1999; MacKillop et al., 2011; Mole et al., 2015; Price et al., 2013). Abnormally high discounting has also been found in a number of psychiatric conditions, including depression, schizophrenia, and bipolar disorder (Amlung et al., 2019). Delay discounting has therefore been proposed as a potential transdiagnostic marker of psychopathology (Amlung et al., 2019; Lempert et al., 2019) and as a risk factor for many health conditions that are largely driven by short-sighted behaviors such as unhealthy diet, smoking, alcohol and drug use (Audrain-McGovern et al., 2009; Fernie et al., 2013; Mozaffarian et al., 2008; Scarborough et al., 2011). The goal of the present study is to identify and validate a predictive fMRI-based brain marker of individual differences in impatience as measured by delay discounting.

A growing number of neuroimaging studies has begun to explore the neurophysiological correlates of individual differences in delay discounting, based on both structural and functional brain measures (Bernhardt et al., 2014; Cooper et al., 2013; Ersner-Hershfield et al., 2009; for a review see Kable and Levy, 2015; Lebreton et al., 2013; Pehlivanova et al., 2018; van den Bos et al., 2014). Several of these studies suggest a role of prefrontal areas, including those involved in reward processing and valuation (especially ventromedial prefrontal cortex/vmPFC and the ventral striatum/VS) (Bartra et al., 2013; Cooper et al., 2013), and those central for cognitive control (especially dorsolateral prefrontal cortex/dlPFC) (Hare et al., 2014). Brain areas associated with memory and prospection, especially hippocampus and entorhinal cortex, are found to contribute to individual differences in delay discounting as well (Benoit et al., 2011; Lebreton et al., 2013; Lempert et al., 2020; Peters and Büchel, 2011). This is in line with findings from psychology and behavioral economics that show that subjective time perception (Zauberman et al., 2009) and the ability to imagine future events (Benoit et al., 2011) and one’s future self (Ersner-Hershfield et al., 2009) in rich detail are contributing to patience and intertemporal decision-making. Several other studies have focused on structural and functional connectivity measures, demonstrating that fiber tract strengths between frontal and striatal areas are associated with decreased discounting, whereas strength of striatal-subcortical connections are associated with increased discounting (van den Bos et al., 2014). The structure of midbrain dopaminergic nuclei and the ventral striatum has also been associated with self-reported trait impulsivity (MacNiven et al., 2020). Thus, previous findings regarding the brain bases of individual differences in impatience and impulsivity offer a mixed picture: different regions seem to be involved and findings are not always replicated across studies.

Given the complexity of intertemporal decisions, it is unlikely that individual differences in delay discounting can be explained by one brain region or one functional network. Instead, they likely result from an interplay between different functional networks, including those associated with the presumed underlying functional processes of valuation, cognitive control, and prospection (Lempert et al., 2019; Peters and Büchel, 2011). However, surprisingly few studies have investigated multivariate or distributed patterns predicting individual differences in delay discounting (Berman et al., 2013; Li et al., 2013; Pehlivanova et al., 2018). Further, most previous studies used relatively small sample sizes for exploring individual differences, increasing the risk of both false-positive and false-negative results (Poldrack et al., 2017). A further limitation of previous findings is that they are often based on correlation or regression models that are not cross-validated on independent (hold-out) data samples (MacNiven et al., 2020). Thus, such models run the risk of overfitting the model to the data, thereby again increasing the risk of false-positive findings that cannot be replicated in independent data sets.

In this paper, we address these limitations by using a machine-learning-based ‘brain model’ approach (Kragel et al., 2018; Woo et al., 2017) in a larger data sample (total N=255). Brain models or ‘brain signatures’ are patterns of activity across the whole brain that are trained to predict a given mental process (e.g., in the case here, impatience as measured by delay discounting) and that can be applied to new, independent data to test their validity and generalizability (Kragel et al., 2018; Poldrack et al., 2017; Scheinost et al., 2019). As such, these models provide a novel approach to analyze fMRI data that goes beyond reporting peak coordinates by providing specific models that can be replicated, validated, or falsified in an easy and quantifiable way. This approach has been successfully applied to brain-based prediction of pain (Wager et al., 2013; Woo et al., 2017), working memory (Rosenberg et al., 2020), affective states (Yu et al., 2020), and physiological measures like heart rate and blood pressure (Eisenbarth et al., 2016; Gianaros et al., 2017), among others.

Here, we build on this approach to predict individual differences in delay discounting. We used an established machine-learning algorithm (Tibshirani, 1996; Wager et al., 2011) on fMRI data from two independent studies, from different scanners, labs, and countries. Study 1 (acquired at Bonn University, N=110) was used for training and cross-validation of a predictive model of individual differences in delay-discounting. Baseline data from Study 2 (University of Pennsylvania, N=145, (Kable et al., 2017)) was used as an independent test dataset to test the validity and replicability of the predictive model. If there is a consistent pattern of functional activity across the whole brain associated with individual differences in delay discounting during intertemporal discounting tasks, then this pattern should be able to predict delay discounting in new data (hold-out subjects) and even completely independent datasets. Further, comparing the resulting pattern to specific meta-analysis-based masks allows us to assess the contribution of brain areas associated with valuation and affect, cognitive control, and memory/prospection.

## Results

### Individual differences in delay discounting

We analyzed behavioral and fMRI data from two independent studies and two different versions of a standard intertemporal choice task (ITC). In Study 1 (collected at Bonn University, N=110 male participants), the ITC task consisted of 108 incentive-compatible choices between smaller sooner (SS) and larger later (LL) monetary rewards. Both SS and LL amounts varied. LL options had delays from 2 to 240 days (see Fig. 1A and Methods for details). Participants’ choices allowed us to compute the discounting parameter *k*. This log(*k*)-parameter describes the steepness of discounting as modeled by the hyperbolic discounting function (Kirby and Herrnstein, 1995). Higher log(*k*) parameters reflect steeper discounting and thus greater impatience, and smaller (more negative) values reflect less steep discounting and thus more patient decision-making.

**Fig 1.**
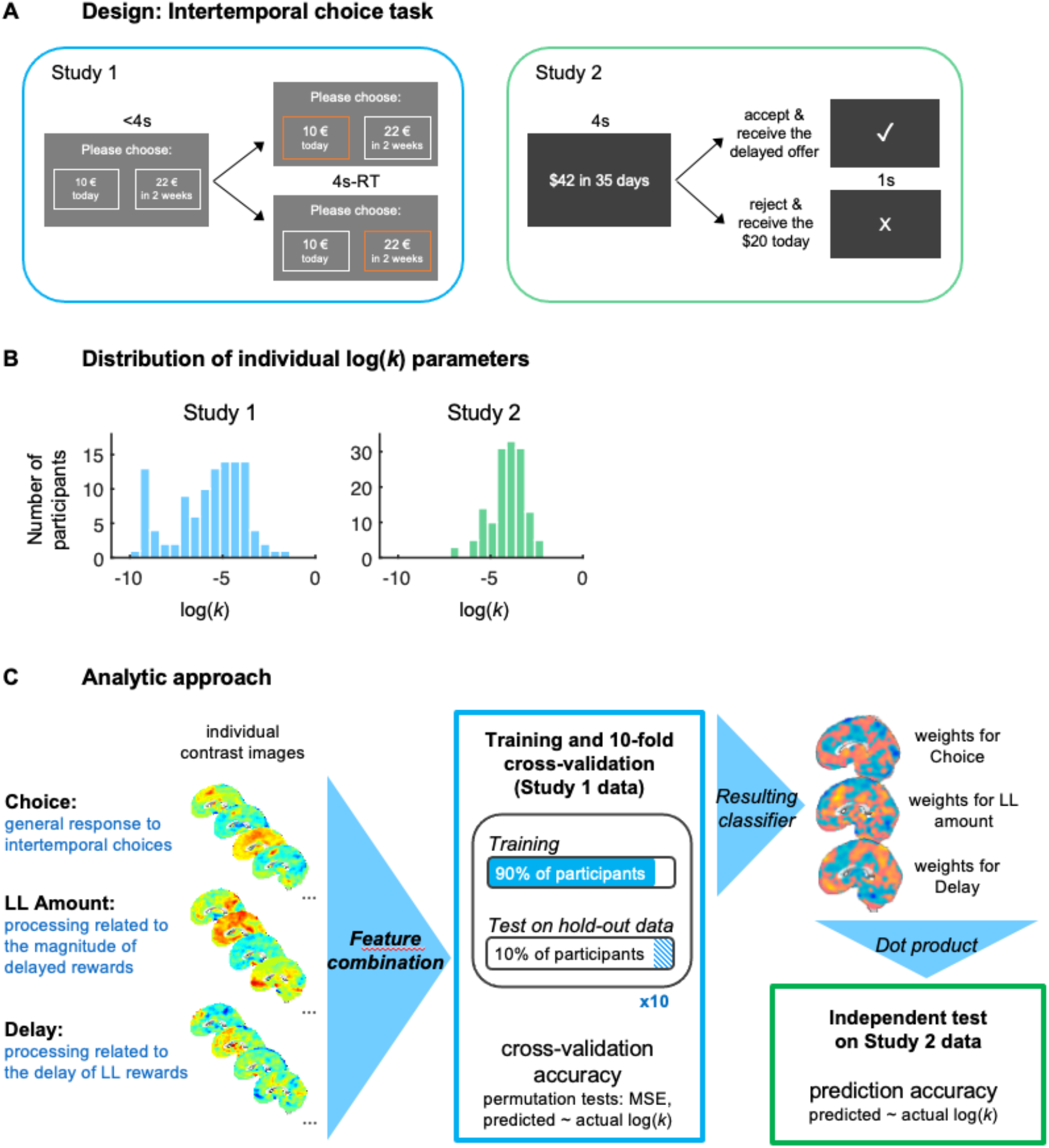
Experimental design, discounting behavior, and analytical approach. **A)** Visual presentation of intertemporal choices (ITC) and their timing in Study 1 and Study 2. **B)** Distribution of individual log-transformed k-parameters by study. **C)** Analytic approach. Contrast images for brain activity in response to the onset of the Choice screen and its parametric modulation by LL Amount and Delay were computed for each participant and concatenated. Data from Study 1 was used for training and 10-fold cross-validation. In each fold, the classifier was trained on 90% of the data using LASSO-PCR and tested on the remaining 10% hold out data to evaluate its predictive accuracy. The predictive classifier obtained from Study 1 was then tested on Study 2 data, acquired on a different scanner, in a different lab and country, assessing its validity in a completely independent dataset.

On average, participants chose the SS option in 43.7% of all trials (median = 48.1%) and had a fitted log(*k*) parameter of -5.70 (median log(*k*) = -5.28, corresponding to a *k* of 0.0051). Choice behavior was characterized by substantial individual differences, with %SS choices ranging from 5.6% to 88.8%, and log(*k*) ranging from -9.90 to -1.36 (see Fig. 1B).

In Study 2 (collected at the University of Pennsylvania, N=145), participants made 120 incentive-compatible choices, in which they were presented varying amounts of LL rewards and delays. Participants had to either accept or reject the LL option, in which case they would receive a fixed SS amount of US$20 (see Fig. 1A). On average, participants in Study 2 chose the SS option in 57.4% of trials (median = 60.0%) and had an average fitted log(*k*) of -4.08 (median log(*k*) = -3.95, corresponding to a *k* of 0.0193). Again, individuals varied substantially in their intertemporal preferences, with %SS ranging from 0.8% to 99.2%, and log(*k*) ranging from -7.08 to -2.12 (see Fig. 1B).

Taken together, our data confirm the substantial variability in impatience known from previous work (Kable and Glimcher, 2007; Pehlivanova et al., 2018). They show that both data sets had sufficient variability in discounting behavior to investigate the neurofunctional bases of these individual differences.

### Significant cross-validated prediction of delay discounting based on fMRI

We applied a LASSO-PCR (least absolute shrinkage and selection operator-principal component regression) (Tibshirani, 1996; Wager et al., 2011)—a machine learning algorithm especially suitable for the multidimensionality of brain imaging data (Wager et al., 2011) to the fMRI data of Study 1. More specifically, we trained a multivariate classifier—a pattern of brain activity across the whole brain (gray-matter-masked)—to reliably predict individual differences in log(*k*). This ‘*k*-marker’ (see Fig. 2A, more details below) was trained based on a feature space that combined individual contrast images for three regressors of a standard univariate GLM first-level model: *Choice* onset, its parametric modulation by *Relative LL Amount*, and its parametric modulation by *Delay* (see Fig. 1C, Methods for details). It captures a combination of functional processes that together determine delay discounting. The predictive accuracy of the classifier (*k*-marker) was assessed using a 10-fold cross-validation procedure in which each participant’s contrast images were included in nine training folds and held out in a tenth fold to assess the *predicted* log(*k*) and thereby the accuracy of the model (see Fig. 1C). This training and cross-validation procedure resulted in a cross-validated prediction-outcome correlation of *r* = 0.49 (permutation test: *p* < 0.0002), a mean squared error of 2.84 (permutation test: *p* < 0.0002), and a mean absolute error for predicted log(*k*) of 1.32 (permutation test: *p* < 0.0002, see Fig. 2B and Suppl. Fig. S1).

**Fig 2.**
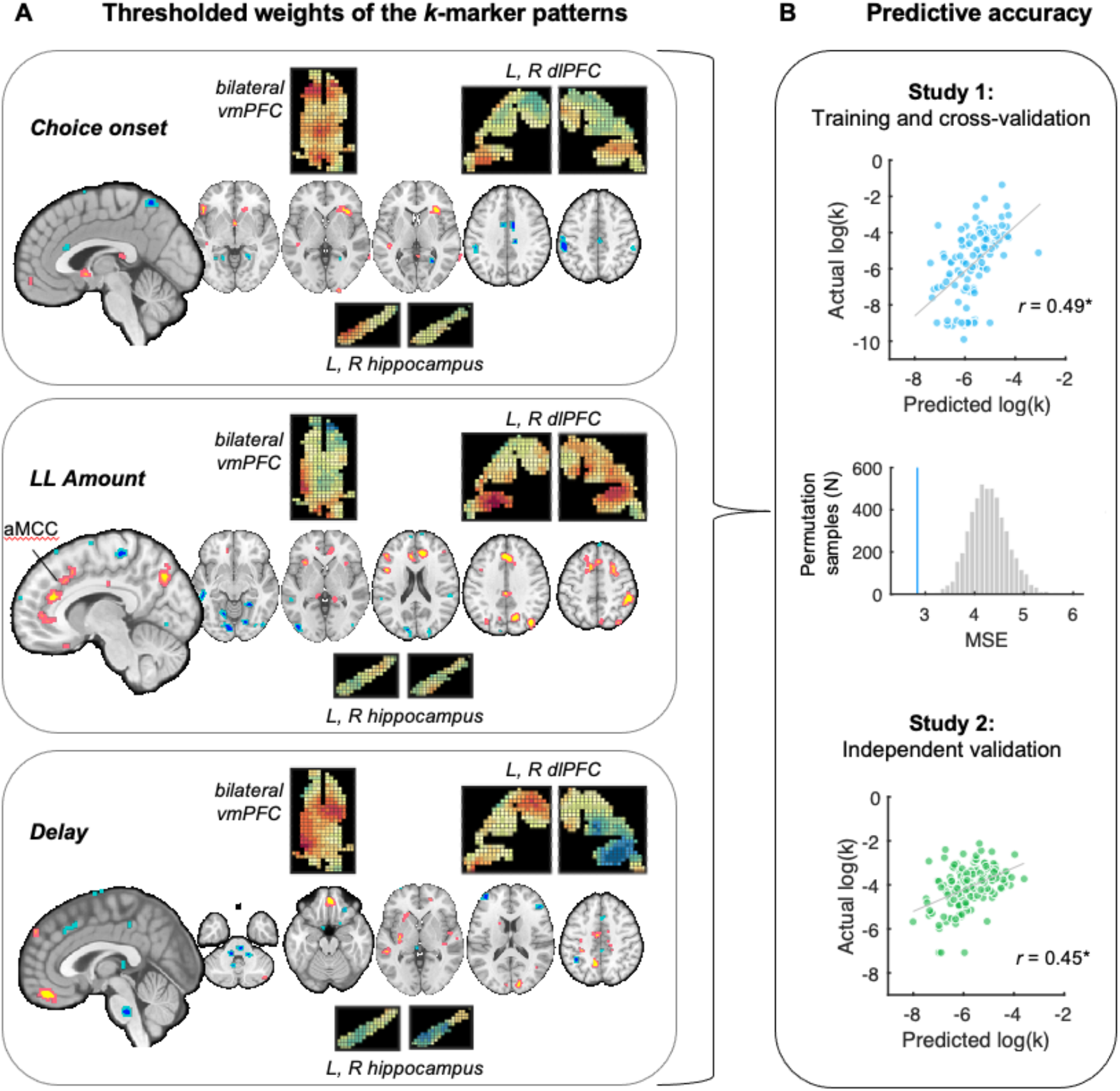
Weight maps and predictive accuracy of the classifier (‘*k*-marker’). **A) Classifier weights for the three contrast images** (Choice onset, parametric modulation by LL Amount and Delay), thresholded for illustration at *q* < 0.05 FDR corrected across the combined feature space (three concatenated maps). Note that *un*thresholded maps are used for prediction and that the combined weights across maps are contributing to the prediction. Pop-out maps show the unthresholded patterns for selected regions of interest (transversal slices for vmPFC, coronal for dlPFC, and saggital for hippocampus), in order to illustrate the heterogeneity of voxel weights (e.g., positive versus negative) within each region and across the three interdependent weight maps. **B) Predictive accuracy**. For interpretability and comparability across both studies, prediction-outcome correlations are shown in scatter plots. Permutation tests (5000 samples) were used to evaluate the significance of the accuracy metrics, mean squared error (MSE, see histogram of null distribution in gray and actual value for Study 1 in blue) and prediction-outcome correlation (see Suppl. Information S1 for additional permutation test results) in Study 1. For Study 2 (independent validation data set), accuracy is assessed as the parametric prediction-outcome correlation. Mean absolute error is provided in the text.

### Validation of the k-marker in an independent test data set (Study 2)

Brain markers of individual differences based on brain images become more meaningful if the results can be applied and validated (i.e., confirmed versus falsified) in different and completely independent datasets (Kragel et al., 2018). The validity of the marker should not depend on study-specific parameters like the type of scanner used for data acquisition, preprocessing software, or other aspects of the dataset (Kragel et al., 2018; Scheinost et al., 2019; Woo et al., 2017).

We therefore tested whether the *k*-marker—developed and cross-validated entirely on Study 1—could be applied to a completely independent dataset. The validation dataset (Study 2) was acquired on a different scanner, in a different lab and country, using a different participant sample and different task characteristics, and was preprocessed and analyzed using a different MRI analysis software. Evaluating the performance of the *k*-marker in Study 2 is therefore an even stronger test of the robustness and replicability of the pattern than cross-validation in Study 1 alone.

For this purpose, we computed the pattern expression of the *k*-marker using the matrix dot product for each participant’s data (contrast images for Choice-onset, parametric modulation for LL-amount and Delay) in Study 2. The resulting predicted log(*k*) values were significantly correlated with actual log(*k*) values (*r* = 0.45, *p* = 1.06*10^−8^, mean absolute error of 1.68) demonstrating the replicability of the *k*-marker in a completely independent dataset.

The training and cross-validation dataset (Study 1) consisted of male participants only, which might limit the validity of the *k*-marker in females. We therefore assessed the accuracy of the *k*-marker in Study 2 separately in male and female participants. In male participants (N=88), the prediction-outcome correlation was *r* = 0.40 (*p* = 1.17*10^−4^). In females (N=57), the prediction-outcome correlation was *r* = 0.55 (*p* = 7.63*10^−6^) and thus very comparable, if not superior, to the predictive accuracy in males (see Suppl. Figure S2). This demonstrates that the *k*-marker (despite being trained on male participants’ data only) predicts delay discounting well for both male and female participants.

### Response of the k-marker differs between lean and overweight participants

Given previous findings of higher delay discounting in overweight and obese people (Amlung et al., 2016; Jarmolowicz et al., 2014), we next tested whether individual expression of the *k*-marker was associated with individual differences in body mass and overweight (as measured by the body mass index [BMI] > 25). Interestingly, neither in Study 1 nor in Study 2 was actual log(*k*) related to BMI. However, in Study 1, response of the *k*-marker significantly correlated with both BMI (*r* = 0.26, *p* = 0.0054, Fig. 3A) and percentage of body fat (*r* = 0.28, *p* = 0.0047). The *k*-marker response in Study 1 also differed between lean (BMI<25) and overweight-to-obese (BMI>25) participants (*t*(108) = 2.85, *p* = 0.0052, Cohen’s *d* = 0.55 see Fig. 3B). In Study 2, the correlation between predicted log(*k*) and BMI was positive but not significant (*r* = 0.09, *p* = 0.28). However, paralleling the findings in Study 1, overweight-to-obese participants had a significantly higher *k*-marker response (higher predicted discounting) compared to lean participants (*t*(143) = 2.11, *p* = 0.037, Cohen’s *d* = 0.35, see Figure 3B).

**Fig 3.**
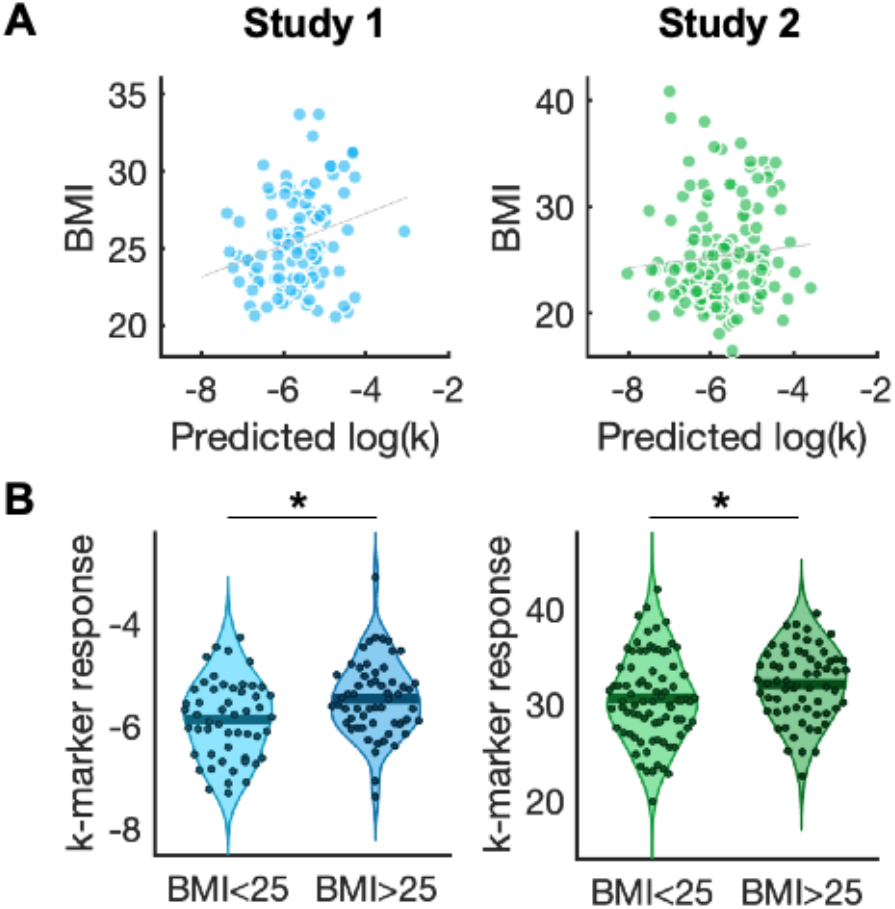
Association of *k*-marker response with BMI and overweight. **A)** In Study 1 (but not Study 2), *k*-marker response was positively and significantly correlated with BMI. **B)** In both studies, *k*-marker response was significantly greater (predicting more delay discounting) in overweight (BMI > 25) compared to lean participants.

### Relationship with other person-level variables

In order to delineate the discriminant and convergent validity of the *k*-marker, we next tested the relationship of the *k*-marker response to several other trait variables that have been associated with delay discounting. Delay discounting could be related to risk preferences because future rewards may be seen as less certain than immediate rewards (but see conflicting evidence, e.g., Green and Myerson, 2013; Rachlin et al., 1991). In Study 1, risk preferences were not significantly associated with *k*-marker response (*r* = - 0.05, *p* = 0.58). However, in Study 2, the response of the *k*-marker was negatively associated with individual risk preferences (alpha estimates, *r* = -0.20, *p* = 0.016). Delay discounting has also been associated with differences in general intelligence (Shamosh et al., 2008). However, the *k*-marker response was not associated to a screening measure for intelligence (assessed in Study 2 only, using the Shipley Institute of Living Scale – Second Edition (Zachary and Shipley, 1986), *r* = 0.06, *p* = 0.44). Delay discounting may also decline with increasing age (Green et al., 1994; Reimers et al., 2009; but see Seaman et al., 2018). While in Study 1 (which had a relatively large age range) there was no significant correlation between age and *k*-marker response (*r* = -0.01, *p* = 0.90), higher age was associated with somewhat lower *k*-marker responses (predictive of less discounting) in Study 2 (*r* = -0.17, *p* = 0.043).

### Thresholded activation patterns of the k-marker

Activation patterns across the whole brain gray matter and across all three contrasts are used for prediction and cross-validation. To illustrate this pattern and to identify the areas that contributed the most strongly with positive or negative weights, we used a bootstrapping procedure (5000 bootstrap samples). Bootstrapped weights were thresholded at *q* < 0.05 FDR corrected across the whole weight map of the combined feature space and displayed in Fig. 2A (see Suppl. Tables S3-5 for full results tables).

Our results revealed a distributed network of areas that jointly contributed to individual differences in delay discounting, including the vmPFC, ventral striatum, anterior midcingulate cortex (aMCC), hippocampus, frontoparietal and visual areas (see Fig. 2). Predictive activity patterns differed for the processes captured by the three different contrast images. Of note, some regions showed negative weights (i.e., predicted less discounting) for one, but positive weights (i.e., predicting more discounting) for another contrast. More specifically, for the Choice (versus implicit baseline) contrast, activity in the striatum, the anterior insula and lateral prefrontal areas contributed positively to more impatience (i.e., greater discounting), whereas activity in bilateral visual areas, premotor and motor areas contributed weights towards less impatience (i.e., less discounting). For the parametric modulation by LL Amount, activity in vmPFC, aMCC, posterior cingulate cortex (PCC), precuneus, and frontoparietal areas were contributing positive weights for greater discounting, whereas visual areas and premotor areas contributed mainly negative weights. For the parametric modulation by Delay, positive weights were observed in the most ventral part of the vmPFC (or mOFC), premotor areas, and visual cortex, while negative weights were observed in frontoparietal areas and several regions of the brainstem.

### Similarity of k-marker brain patterns to meta-analytic maps and established functional networks

To investigate the similarity of these predictive patterns of the *k*-marker with functional networks, we next compared them with term-based meta-analytic images (Yarkoni et al., 2011) for terms that may contribute to intertemporal decision-making (**Fig. 4**) and canonical resting-state networks (Yeo et al., 2011) (**Fig. S3**). We thus computed the spatial correlation (Pearson’s *r*) between the *k*-marker and these specific maps of interest (c.f., Koban et al., 2019). While these correlations are descriptive, they can inform us quantitatively whether and in which direction (positive or negative) previously identified functional networks contribute to individual differences in delay discounting. They can also hint at the functional processes that may underlie the *k*-marker patterns.

**Fig 4.**
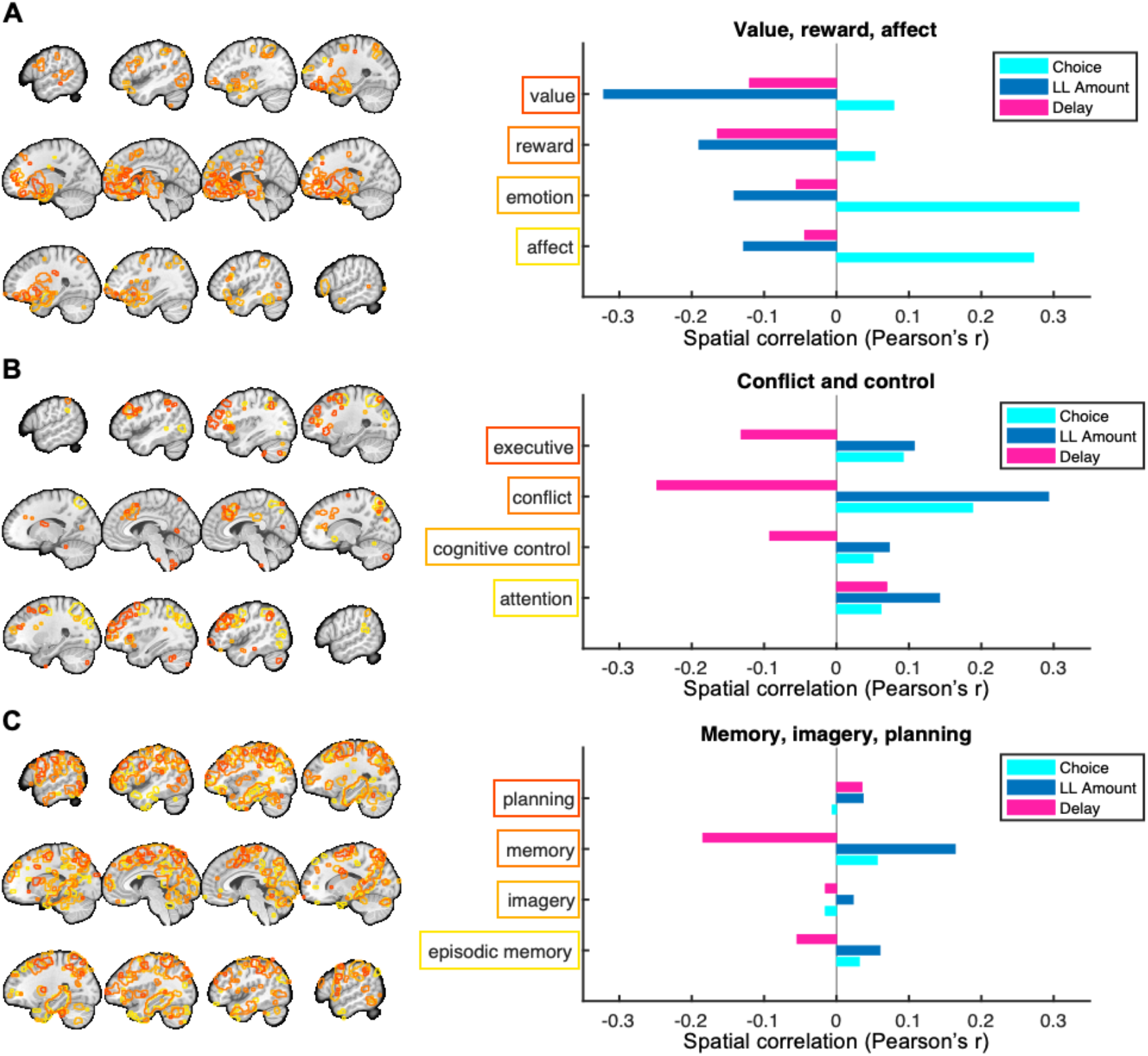
Spatial similarity of the *k-marker* with meta-analytic maps. In order to quantitatively compare the classifier patterns with theoretically relevant functional networks, we computed the spatial correlation of the *k*-marker patterns with Neurosynth meta-analytic maps (Yarkoni et al., 2011) associated with **A) value and affect, B) conflict and cognitive control, and C) memory and prospection**. Meta-analytic maps from each group of terms are overlaid on the left (outline colors matching the outlines of the terms on the right) and can be inspected in greater detail online (www.Neurosynth.org). Note that spatial correlations are purely descriptive, indicating whether activity in any of the shown functional maps is rather positively (*r* > 0) or negatively (*r* < 0) associated with discounting, for each component map of the *k*-marker (Choice, LL Amount, Delay).

We compared the similarity of the *k*-marker with meta-analytic maps for 1) affective and value-related, 2) conflict and cognitive-control related, and 3) memory-related processes (see **Fig. 4**). Value- and affect-related maps (especially ‘affect’ and ‘emotion’) showed consistent positive correlations (*r’s* > 0.05) with the Choice-related pattern of the *k*-marker, in line with the idea that more affect-related activity during intertemporal choices overall leads to more *impatient* decisions (greater discounting). However, stronger engagement of affective and especially ‘reward and ‘value’-related activity for increasing LL-Amount (and, to a lesser extent, for increasing delays) was associated with less discounting (i.e., more patience). This suggests that greater patience is associated with greater sensitivity of valuation-related signals to the amount of LL rewards.

In contrast to our initial hypothesis, cognitive control related activity was not strongly or even negatively associated with more impatience (greater discounting). Instead, there were *positive* correlations (*r*’s from 0.05 to 0.19) of the Choice-pattern with meta-analytic maps for the terms ‘attention’, ‘cognitive control’, ‘conflict’, and ‘executive’ (more positive weights predicting greater discounting). Further, stronger activation of control-related maps by greater LL-Amount was associated with greater discounting (more impatience), whereas stronger activation of control-related activity for longer delays was associated with less discounting (more patience). Parallel findings were observed for the frontoparietal network, which is tightly associated with cognitive control (see Suppl. Materials and **Fig. S3**). This suggests that control-related activity is not directly and positively related to patience, but that it is engaged in more nuanced ways such as processing amounts and delays.

Finally, we assessed the contribution of brain systems related to memory or prospection. While the term ‘memory’ (which likely also includes many working-memory studies) showed a similar pattern as control-related maps, more specific terms such as ‘episodic memory’, ‘imagery’, or ‘planning’ were not substantially positively or negatively correlated with any of the *k*-marker patterns (*r*’s around 0.05 and smaller).

## Discussion

A major goal of neuroscience and psychiatry is to identify brain markers of transdiagnostic processes that may be altered across different diseases or that predispose individuals to such diseases (Insel and Cuthbert, 2015). Impatience, measured by how much people discount future compared to immediate rewards, has been proposed as such a transdiagnostic process across psychopathology, addiction, obesity and eating disorders (Amlung et al., 2019; Bickel et al., 2014; Lempert et al., 2019). In this paper, we aimed to advance our understanding of the brain processes that drive variability in impatience by identifying and validating a distributed pattern of functional brain activity that predicts individual differences in delay discounting. We first used a cross-validation procedure to develop a novel brain-marker of delay discounting (*k*-marker) based on whole brain, grey-matter masked fMRI data (N_1_=110). We then validated the *k*-marker in an independent second fMRI data set (N_2_=145) sampled in a different cohort with different socio-demographic characteristics, a different fMRI scanner, and employing a different delay discounting task. An important advance of the present study is that it provides a novel ‘brain model’ (Kragel et al., 2018) of impatient decision-making. As such, this brain model can be further tested, validated, or refined in future studies, including those that test its generalizability and predictive power in clinical populations.

Our results also inform the debate regarding the contributions of specific brain regions and functional networks to understand people’s ability to delay gratification. Among the brain areas that contributed most strongly with positive and/or negative weights were the vmPFC, striatum, and other regions associated with valuation and reward (Bartra et al., 2013; Clithero and Rangel, 2014; Levy and Glimcher, 2011). This finding is in line with previous, univariate findings (Cooper et al., 2013; MacNiven et al., 2020; Pehlivanova et al., 2018). The present results add to this emergent picture, by suggesting that overall reward sensitivity is associated with greater discounting, whereas greater sensitivity to the amount of the reward is linked to more patient decision-making.

Significant weights were also found in the frontoparietal control network, which is associated with cognitive control, and has been proposed to implement self-control and far-sighted, patient decision-making (Hare et al., 2014; McClure et al., 2004). Yet, the present results are surprising as they draw a more complex picture of these areas’ contribution to delay discounting, with modulation of these areas by greater LL rewards being positively associated with impatience, and modulation by delay being negatively associated with impatience. Thus, cognitive control-related areas were more recruited for long delays and small LL amounts for low discounters, and shorter delays and larger LL amounts for high discounters. These are the cases in which decisions are most difficult (closer to the indifference point) and therefore require resolution of response conflict (Botvinick et al., 2001; Hutcherson et al., 2020; Kool et al., 2013; Shenhav et al., 2013). These findings have at least three important implications for models of delay discounting and self-control in cognitive neuroscience. First, they speak against the idea of a simple dual process account of intertemporal choice and self-control (McClure et al., 2004), joining previous work that suggested more complex neural processes at play (Ballard and Knutson, 2009; Berkman et al., 2017; Hare et al., 2014; Kable and Glimcher, 2007). Second, it also speaks against the idea that more frontoparietal activity is related to higher individual levels of self-control. Instead, it suggests that *for which choice options* control-related areas are activated is more informative than their overall level of activation. This finding is in line with value-based choice models of self-control (Berkman et al., 2017) and with recent evidence that high and low discounters differ in how much attention they allocate to amount versus delay information (Amasino et al., 2019). It also fits with the idea that low discounters may not need ‘control’ to discount less (Lempert et al., 2019), and that high discounters may employ cognitive control for different types of decisions. Third, our results thereby also suggest a mostly neglected role for conflict detection in inter-temporal choice feeding into cognitive control mechanisms.

Hippocampus and adjacent midtemporal areas have been associated with individual differences in discounting (Benoit et al., 2011; Lebreton et al., 2013; Peters and Büchel, 2011), in line with the importance of prospection and self-projection in intertemporal decision-making. While our pattern shows some weights in parahippocampal areas and in occipital areas, these weights seemed to be less strong than those in other brain regions. The *k*-marker patterns were not clearly associated with meta-analysis-based activation maps of episodic memory or prospection. This suggests that, at least in the present data of neurotypical healthy adults, activation in these areas might be less related to individual differences than activation patterns in prefrontal and parietal ones. Yet, this does not preclude that interactions between hippocampus and prefrontal areas might be important or that damage in these regions may lead to substantial changes in delay discounting. Together, our findings highlight the importance of a distributed activity pattern across the whole brain instead of single regions, confirming the notion that impatience and delay discounting depend on the interactions between different functional processes and networks in the brain.

Previous work has related individual differences in delay discounting with obesity, substance use disorders, and multiple psychiatric diseases (Amlung et al., 2019; Lempert et al., 2019; Peters and Büchel, 2011). In the two samples presented here, log(*k*) values based on participants choices themselves were not significantly associated with BMI or overweight. Important to note, the two studies were not directly designed to include a large range of BMI or a high number of obese participants, which may explain this null effect. However, responses to the *k*-marker did significantly differ between lean and overweight participants. This finding suggests that this brain marker is reflecting variance in functional neurophysiology that is related to long-term patterns in decision-making and health parameters. It also may imply that the brain marker is more sensitive than the behavioral measures it was trained on to speak to individual differences in delay discounting. We call for future studies to test the *k*-marker in samples with larger numbers of participants with obesity to further investigate this idea. Future studies could also assess whether the *k*-marker has prognostic value for weight-related health outcomes such as cardiovascular disease or diabetes and responsiveness to drastic weight loss interventions such as bariatric surgery.

The present study also provides some initial evidence for how specific the *k*-marker is to delay discounting, by dissociating it from other person-level variables that are often associated with the degree of impatience. For example, responses of the *k*-marker were not (Study 1) or only moderately (Study 2) associated with risk preferences or participants’ age. They also did not correlate with a simple measure of general intelligence (only assessed in Study 2). These results support the interpretation that the *k*-marker is indeed a measure of discounting, rather than reflecting other, potentially confounded variables. Future work could more closely examine the relationship of the *k*-marker with other related mental processes, with personality, and with clinical measures of interest.

The purpose of the present study was to investigate the functional brain activation during intertemporal choices that is predictive of individual differences in delay discounting. As a resulting limitation, the *k*-marker is linked to specific task-related contrasts. It cannot be readily applied to task-independent data such as resting-state fMRI or structural MRI that might be more readily available in existing clinical or large-scale datasets. Future studies could use a similar machine-learning approach to develop markers of delay discounting based on task-independent brain measures. They could also extend this work by investigating the role of structural and functional connectivity and the potential contribution of structure-function interactions to individual differences in delay discounting.

A strength of the *k*-marker developed in this paper is that it can be applied to and tested in any other fMRI study on monetary delay discounting, for which contrast images for Choice, LL Amount, and Delay can be computed. Future work could also test the validity of the *k*-marker in other types of delay discounting paradigms that involve non-monetary rewards such as food, and to other, similar paradigms such as social discounting (Jones and Rachlin, 2006; Strombach et al., 2015).

The classifier (*k*-marker) predicts these individual differences in neurotypical healthy adults across different populations, scanners, and analysis pipelines. Future work could test the generalizability of the *k*-marker in children, adolescents, the elderly, or in clinical populations. Such work could assess whether the *k*-marker is able to predict clinical status and health outcomes in conditions related to abnormal discounting, such as eating disorders, substance use, and other psychiatric disorders (Amlung et al., 2019; Lempert et al., 2019).

## Methods

### Participants

For Study 1, participants were recruited in the context of a longitudinal dietary intervention study at the University of Bonn, Germany (https://osf.io/rj8sw/?view_only=af9cba7f84064e61b29757f768a8d3bf). Due to the nature of this intervention study, we only recruited male participants who further fulfilled the following inclusion criteria: age between 20-60 years, right-handedness, non-smoker, no excessive drug or alcohol use in the past year, no psychiatric or neurological disease, body mass index (BMI) between 20-34, no other chronic illness or medication, following a typical Western diet without dietary restrictions, no MRI exclusion criteria (large tattoos, non-removable piercings, metal in the body, claustrophobia, etc.). N=116 male participants performed the ITC task in Study 1. Data reported here was collected during a baseline session before the group assignment and dietary intervention (to be reported elsewhere). The data of 6 participants had to be excluded for analysis due to following reasons: technical problems with the scanner (1), with the synchronization between stimulation software and scanner (3), with the response box (1), and due to strong motion artefacts (>5mm) and participant quitting the task mid-scan (1). Therefore, 110 participants (mean age = 31.7y) were included in the final analysis of Study 1.

Data from Study 2 was conducted in the context of a large cognitive training study at the University of Pennsylvania, PA, USA (Kable et al., 2017). The goal was to examine if commercial cognitive training software leads to significant changes in decision-making behaviors including delay discounting. Participants completed two sessions of scans 10 weeks apart. Here we only include the baseline (pre-intervention) data. Of the 160 non-pilot participants that completed session 1, we excluded those with missing runs (n = 6), frequent or significant head movement (any run with >5% of mean image displacements greater than 0.5mm; n = 3), more than 3 missing trials per run for two or more runs (n = 2), or lack of participant blinding (n = 1, one subject expressed awareness of their experimental condition, i.e. cognitive training vs. control). Of the remaining 148, we excluded three more participants whose choice was entirely one-sided (i.e., choosing only immediate reward or delayed reward), resulting in a final sample of N = 145 participants for Study 2 (88 male, 57 female, mean age = 24.4y).

All participants were paid for their participation in the study. The study protocols were approved by the IRB of Bonn University’s Medical School (Study 1) and the University of Pennsylvania (Study 2).

### Stimuli and task

In Study 1, participants performed 108 choices (trials) between varying amounts of smaller sooner (SS) and larger later (LL) options, presented on the left or right of the screen (position randomized, see Fig 1A). Participants were instructed that one of their choices might be paid out at the end of the experiment. Thus, participants’ choices were non-hypothetical and incentive-compatible. During each trial, the two options were presented for 4s, during which participants could make their choice (left or right) by pressing the corresponding response key with their left or right index finger, respectively. Once the choice has been made, a yellow frame highlighted the chosen option and remained on the screen for the remainder of the 4s. Inter trial intervals were jittered using an approximately geometric distribution (2-11s)

SS options varied between 5, 10, and 20 Euro, and always had zero delay (‘today’). LL options varied between 5 and 96.80 Euro and had delays between 2 days and 8 months (∼240 days). Amounts and delays were chosen in order to allow fine-grained estimation of individual *k*’s between 0 and 0.256. Suppl. Table 1 specifies all combinations of SS, LL, delays, and corresponding indifference *k*’s (*k*’s at which the LL and SS options would be chose at equal proportion). Trials were presented in randomized order.

The ITC task in Study 2 consisted of 120 trials, with again the same choice sets for all participants. In contrast to Study 1, SS amount was fixed to US$20. Thus, participants were only presented with the LL option (with amount ranging from US$22 to US$85, and delays from 19 days to 180 days) and were instructed to press one of two keys to either accept and receive this LL offer, or to reject the LL offer and receive the SS offer ($20 today) instead. Participants were informed that one trial would be randomly chosen at the end of the experiment and participant’s choices implemented, resulting in incentive-compatible and mutually independent choices in each trial (as in Study 1).

## Data analyses

### Behavioral measures

For each participant, we calculated the proportion of SS choices (with respect to total number of non-misses) and the model-based discounting parameter *k*. More specifically, in Study 1 we computed *k* by calculating the proportion of SS choices for all *target k*’s (i.e., the k-value for which SS and LL options of any given choice trial should theoretically be chosen at 50% each). We then used linear interpolation to identify the individual indifference point at which the proportion of SS and LL choices was equal (50% each).

In Study 2, we fit a logit utility model on choice data via maximum likelihood estimation. The logit of the probability of choosing the delayed reward was modeled as follows:

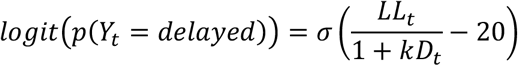

Where *LL*_*t*_ is the LL amount in trial t and *D*_*t*_ is the delay in trial t. σ was included as a scaling parameter that controls the relationship between utility difference scale and choice. Individual *k*’s were log-transformed in both studies to obtain non-skewed distributions of discounting parameters.

### MRI data acquisition

Functional and structural brain imaging data for Study 1 was acquired using a Siemens Trio 3T scanner (Erlangen, Germany) at the Life & Brain Institute, Bonn University Hospital, Germany. Functional images used a T2* weighted EPI-GRAPPA sequence (TR = 2.5s, TE = 30ms, flip angle = 90**°**, FOV = 192mm) and covered the whole brain in 37 slices (voxel size of 2×2×3mm). Structural images were acquired using a T1 weighted MPRAGE sequence (1mm isomorphic voxels).

For Study 2, the functional and structural imaging data was acquired with a Siemens 3T Trio scanner with a 32-channel head coil. High-resolution T1-weighted anatomical images were acquired using an MPRAGE sequence (T1 = 1100ms; 160 axial slices, 0.9375 x 0.9375 x 1.000 mm; 192 x 256 matrix). T2*-weighted functional images were acquired using an EPI sequence with 3mm isotropic voxels, 64 x 64 matrix, TR = 3,000ms, TE = 25ms, 53 axial slices, 104 volumes. B0 fieldmap images were collected for distortion correction (TR = 1270ms, TE = 5 and 7.46ms).

### Preprocessing and basic statistical analyses of fMRI data

Preprocessing for Study 1 was performed in SPM12 and used a standard pipeline of motion correction, slice time correction, spatial normalization to MNI space, and spatial smoothing of images using an 8mm FWHM Gaussian kernel. Preprocessing for Study 2 was performed in FSL according to the original preprocessing pipeline for Kable et al. (2017). This involved the standard pipeline of motion correction, b0 map unwarping, interleaved slice time correction, spatial smoothing with FWHM 9mm Gaussian kernel, and high-pass filtering (cutoff = 104s).

For Study 1, we used SPM12 to fit a general linear model (GLM) for each participant’s imaging data, with choice onset modeled as a stick function (0s duration) as the main regressor and mean-centered parametric modulators for delay, relative LL amount, SS amount, and reaction time. Six nuisance regressors were added to control for movement artefacts.

For Study 2, FSL was used to fit an otherwise similar statistical model with a choice onset as main regressor and mean-centered parametric modulators for delay, LL amount, and reaction time. As in Study 1, six movement regressors were added to control for head movement.

Individual contrast images were calculated for the following three regressors of interest that were available for both studies: 1) Choice onset versus implicit baseline, 2) parametric modulation by (relative) LL amount, and 3) parametric modulation by Delay. Contrast images were gray-matter-masked, in order to remove voxels that are unlikely to contain meaningful BOLD signal, and individually z-scored to remove differences in scale across images and thereby make results transferable across studies and datasets.

### Training and cross-validation

Training and cross-validation was performed on data from Study 1 only. In order to combine information across all three contrasts images of interest (Choice onset, relative LL amount, and Delay), we concatenated the contrast images for each participant, resulting in a feature space that was triple the size of a single brain image. LASSO-PCR (least absolute shrinkage and selection operator-principal component regression) (Tibshirani, 1996)—a machine-learning based regression algorithm—was then used to train a classifier to predict log(*k*) across all voxel weights of the concatenated contrast images. LASSO-PCR first performs a data reduction using principal component regression, thus identifying brain regions and networks that are highly correlated with each other. It then performs the LASSO algorithm, which shrinks regression weights towards zero, thus reducing the contribution of less important and more unstable components. LASSO-PCR has been shown to be advantageous for brain images for several reasons (see Wager et al., 2011; Wager et al., 2013): it is adequate for predictions based on hundred-thousand voxels, it takes into account multicollinearity between voxels and brain regions, and it yields interpretable results by allowing reconstruction of voxel weight maps based on PCR results.

To assess the predictive accuracy of the classifier in new subjects, we used 10-fold cross-validation. Thus, the training data was split up in 10 stratified combinations of training (90%) and test sets (10%), such that every subject’s data was used for the training of the classifier in 9 folds and held out in the remaining fold to independently assess the prediction-outcome correlation. Tenfold cross-validation was chosen *a priori* as a good compromise between maximizing the sample size in each training set, while at the same time avoiding leave-one-out-cross-validation, which can underestimate accuracy by inducing a systematic bias towards negative prediction-outcome correlations (Poldrack et al., 2020; Scheinost et al., 2019). A priori set default regularization parameters were used for all machine-learning analysis to avoid biasing the model parameters to the data and thereby generating over-optimistic accuracy scores. Permutation tests (5000 iterations of randomly permuting the log(*k*)-values) were used to generate null-distributions and to assess the statistical significance of the prediction-outcome correlation and the mean absolute error.

### Bootstrapping and thresholding

To identify the brain areas contributing with the most reliable positive or negative weights, we performed a bootstrap analysis. 5000 samples with replacements were taken from the paired brain and outcome data and the LASSO-PCR was repeated for each bootstrap sample. Two-tailed, uncorrected p-values were calculated for each voxel based on the proportion of weights above or below zero (Wager et al., 2011; Wager et al., 2013). False Discovery Rate (FDR) correction was applied to p-values to correct for multiple comparisons across the whole feature space (three combined brain maps).

### Independent test set

Study 2 was used as an independent test set to assess validity and generalizability of the brain pattern classifier developed based on Study 1 (i.e., the ‘k-marker’). To assess the response of the predictive marker in Study 2 we calculated the matrix dot product between the *k*-marker and the concatenated contrast images (Choice onset, LL amount, and Delay) from each participant. The dot product reflects the pattern similarity between the classifier and each participant’s set of contrast images and, in sum with the classifier’s intercepts provides a predicted value of *k*. Predictive accuracy of the marker was quantified by correlating the predicted value of *k* with the actual *k*’s of each participant and by calculating the mean absolute error for each prediction.

## Author note

We received funding from the ANR (Tremplin-ERC grant “Brain Gut Decision” to HP), Campus France (Marie-Sklodowska-Curie co-fund fellowship PRESTIGE-2018-2-0023 to LK), French federal funding (program ‘*Investissements d’avenir’*, ANR-10-IAIHU-06), the Federal Ministry of Education and Research Germany (Diet-Body-Brain grant 01EA1809B to BW), the National Cancer Institute (R35CA197461[C.L.] and R01CA170297 [J.K and C.L]). We thank P. Trautner for help with the programming of the task (Study 1), A. Simonetti, M. Boerth, A. Koehlmoos, S. Winkler, L. Bernardo, A. M. Burke, M. K. Caulfield, N. Cooper, G. Donnay, M. Falcone, J. Jorgensen, R. Kazinka, J. Luery, M. McConnell, Rickie Miglin, D. Mukherjee, T. Parthasarathi, S. Price, M. Schlussel, R. Sharp, H. J. Sohn, D. Spence, and K. Terilli for help with data acquisition and T. D. Wager for the Canlab toolbox and helpful discussion.

## Author contributions

LK and HP conceptualized the study. DS acquired the data for Study 1, JK and SL contributed and re-analyzed existing data (Study 2). LK analyzed and visualized the data. LK wrote the first draft of the manuscript, with revisions from HP. HP, JK, CL, MCS, and BW obtained funding, provided resources, and supervised the project. All authors edited and approved the manuscript.

