## Supplemental Materials for "An fMRI-based brain marker predicts individual differences in delay discounting"

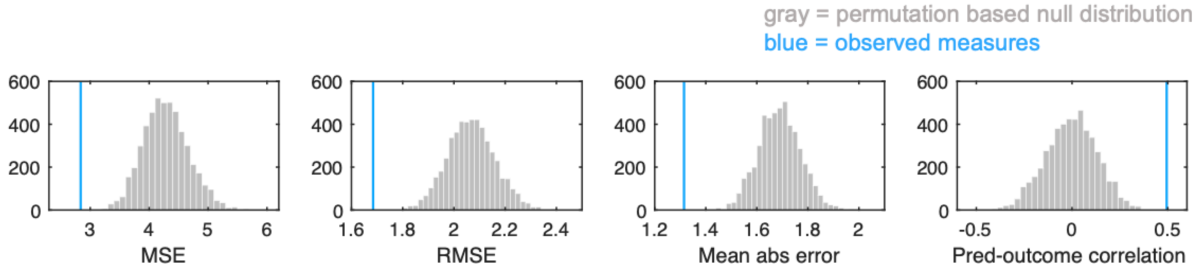


**Fig. S1. Results of the permutation test (Study 1).** Log(*k*) values were randomly permuted, and the prediction algorithm was repeated on the permuted brain-outcome data 5000 times in order to generate null distributions for standard accuracy metrics (from left to right): mean squared error (MSE), root mean squared error (RMSE), mean absolute error (mean abs error), and prediction-outcome correlation (for additional interpretability). Histograms of null distribution are shown in gray bars, observed metrics in blue. For all metrics, observed values were outside the range of permutation samples (all $p$'s $< 0.002$).

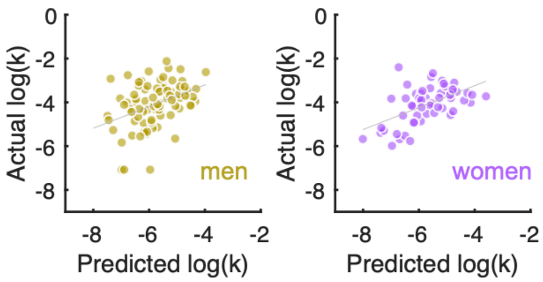


**Fig. S2. Correlations between predicted and observed log(k) were significant in both male and female participants (Study 2).**

**Suppl. Table S3.** Significant positive and negative weights contributing to the *k*-marker during CHOICE (FDR corrected $q < 0.05$ across the whole feature space, i.e. three concatenated gray-matter-masked whole brain maps, and at least three contingent voxels). In the column "Atlas label", cortical regions (Ctx) are labeled based on the multimodal cortical parcellation by Glasser et al. (2016), basal ganglia regions are based on Pauli et al. (2016), cerebellar regions on Diedrichsen, et al. (2009), and brainstem regions based on a combination of studies (Bär et al., 2016; Shen et al., 2013). The entire combined anatomical atlas is available on Github: https://github.com/canlab/Neuroimaging_Pattern_Masks/tree/master/Atlases_and_parcellations/2018_Wager_combined_atlas. This repository includes multiple atlases and other meta-analytic and multivariate maps. Tools for manipulating and analyzing this and other atlases are in the CANlab Core Tools repository: https://github.com/canlab/CanlabCore.

***Positive weights***

| Name | Atlas label | Volume (voxels) | MNI Coordinates X | Y | Z | max(z) |
|---|---|---|---|---|---|---|
| Orbitofrontal cortex | Ctx_a47r_L | 29 | -34 | 40 | -12 | 4.49 |
| Anterior insula | Ctx_FOP5_R | 114 | 32 | 24 | 4 | 4.45 |
| Midtemporal gyrus | Ctx_PHT_R | 18 | 68 | -48 | 4 | 4.43 |
| Frontal pole/vmpFC | Ctx_10pp_L | 29 | -8 | 60 | -12 | 4.42 |
| Midtemporal gyrus | Ctx_TE1a_R | 6 | 66 | -4 | -24 | 4.24 |
| Temporal operculum | Ctx_LBelt_L | 32 | -42 | -28 | 2 | 4.21 |
| Other | No_label | 9 | -2 | -28 | 14 | 4.13 |
| Visual cortex | Ctx_V2_R | 18 | 20 | -100 | 0 | 4.04 |
| Orbitofrontal cortex | Ctx_a47r_L | 11 | -44 | 38 | -12 | 4.03 |
| Ventrolateral prefrontal cortex | Ctx_45_L | 72 | -50 | 28 | -2 | 4.03 |
| Striatum | V_Striatum_R | 38 | 18 | 26 | 0 | 3.99 |
| Striatum | V_Striatum_L | 22 | 0 | 4 | -2 | 3.96 |
| Orbitofrontal cortex | Ctx_11l_L | 13 | -18 | 42 | -14 | 3.95 |
| Retrosplenial cortex | Ctx_RSC_R | 15 | 2 | -46 | 8 | 3.94 |
| Temporal operculum | Ctx_PoI1_L | 20 | -38 | -4 | -16 | 3.89 |
| Visual cortex | Ctx_V4_R | 3 | 30 | -92 | 22 | 3.86 |
| Other | No_label | 3 | 66 | -54 | 8 | 3.84 |
| Visual cortex | Ctx_V3_L | 3 | -22 | -94 | 24 | 3.82 |
| Midtemporal gyrus | Ctx_TE1a_L | 14 | -64 | -10 | -14 | 3.79 |
| Amygdala | Ctx_PeEc_R | 5 | 20 | 6 | -26 | 3.75 |
| Caudate | Cau_L | 3 | -26 | -32 | 20 | 3.72 |
| Precentral gyrus | Ctx_4_L | 9 | -12 | -26 | 70 | 3.71 |
| Temporal pole/amygdala | Ctx_TGd_R | 7 | 34 | 12 | -24 | 3.65 |
| Visual cortex | Ctx_V2_L | 4 | -14 | -102 | 14 | 3.62 |
| Midtemporal gyrus | Ctx_TGd_R | 4 | 52 | 4 | -26 | 3.59 |
| Orbitofrontal cortex | Ctx_13l_L | 4 | -28 | 28 | -14 | 3.48 |

***Negative weights***

| Name | Atlas label | Volume (voxels) | X | Y | Z | max(z) |
|---|---|---|---|---|---|---|
| Midcingulate cortex | Ctx_24dv_R | 58 | 8 | 0 | 36 | -5.03 |
| Cerebellum | Cblm_CrusII_L | 130 | -24 | -76 | -40 | -4.87 |
| Superior frontal gyrus | Ctx_SFL_L | 106 | -8 | 10 | 70 | -4.83 |
| Visual cortex | Ctx_ProS_R | 79 | 24 | -52 | 4 | -4.81 |

| | | | | | | |
|---|---|---|---|---|---|---|
| Inferior parietal lobule | Ctx_PF_L | 338 | -54 | -30 | 42 | -4.75 |
| Precuneus | Ctx_7Am_L | 67 | -2 | -56 | 60 | -4.63 |
| Posterior cingulate cortex | Ctx_23c_R | 51 | 8 | -24 | 40 | -4.35 |
| Parietal cortex | Ctx_PGs_R | 17 | 38 | -70 | 52 | -4.28 |
| Parahippocampal cortex | Ctx_VMV2_L | 7 | -32 | -50 | -6 | -4.06 |
| Midcingulate cortex | Ctx_p24pr_L | 20 | -6 | 2 | 36 | -4.00 |
| Parahippocampal cortex | Ctx_ProS_L | 13 | -20 | -48 | -2 | -3.99 |
| Intraparietal sulcus | Ctx_7PC_L | 11 | -44 | -42 | 60 | -3.94 |
| Cerebellum | Cblm_CrusI_R | 6 | 50 | -48 | -28 | -3.88 |
| vmPFC | Ctx_9m_L | 8 | -8 | 50 | 16 | -3.87 |
| Cerebellum | Cblm_CrusI_L | 6 | -46 | -64 | -42 | -3.87 |
| Midcingulate cortex | Ctx_33pr_L | 6 | -2 | 22 | 20 | -3.84 |
| Inferior parietal lobule | Ctx_PF_R | 13 | 54 | -34 | 46 | -3.83 |
| Inferior frontal cortex | Ctx_6r_L | 3 | -54 | 4 | 16 | -3.80 |
| Dorsolateral prefrontal cortex | Ctx_9_46d_R | 4 | 20 | 48 | 26 | -3.68 |

**Suppl. Table S4.** Significant positive and negative weights contributing to the k-marker for the parametric modulation by LL AMOUNT (FDR corrected $q < 0.05$ across the whole feature space, i.e. three concatenated gray-matter-masked whole brain maps, at least three contingent voxels).

***Positive weights***

| Name | Atlas label | Volume (voxels) | MNI Coordinates X | Y | Z | max(z) |
|---|---|---|---|---|---|---|
| Medial frontal gyrus | Ctx_8BM_R | 386 | -2 | 26 | 36 | 5.86 |
| Dorsolateral prefrontal cortex | Ctx_6a_R | 199 | 26 | 8 | 52 | 5.80 |
| Intraparietal Sulcus | Ctx_PFm_R | 288 | 48 | -36 | 52 | 5.72 |
| Orbitofrontal cortex | Ctx_11l_R | 50 | 16 | 30 | -24 | 5.55 |
| Inferior parietal lobule | Ctx_PGs_R | 298 | 38 | -74 | 36 | 5.40 |
| Dorsolateral prefrontal cortex | Ctx_SCEF_L | 131 | -14 | 14 | 52 | 5.12 |
| Precuneus | Ctx_POS2_R | 177 | 12 | -66 | 38 | 5.12 |
| Cingulate cortex | Ctx_d32_R | 114 | 10 | 34 | 22 | 4.93 |
| Inferior frontal cortex | Ctx_IFJa_L | 134 | -44 | 8 | 26 | 4.88 |
| Ventromedial prefrontal cortex | Ctx_p32_L | 24 | -12 | 44 | 4 | 4.80 |
| Cerebellum | Cblm_IX_R | 18 | 4 | -50 | -34 | 4.75 |
| Orbitofrontal cortex | Ctx_11l_L | 37 | -24 | 44 | -18 | 4.66 |
| Dorsolateral prefrontal cortex | Ctx_p9_46v_L | 67 | -46 | 28 | 22 | 4.66 |
| Visual cortex | Ctx_V1_R | 29 | 18 | -76 | 12 | 4.66 |
| Intraparietal Sulcus | Ctx_AIP_L | 24 | -38 | -44 | 42 | 4.61 |
| Inferior parietal lobule | Ctx_IP1_L | 58 | -32 | -72 | 32 | 4.57 |
| Posterior cingulate cortex | Ctx_23d_R | 56 | -2 | -28 | 36 | 4.56 |
| Anterior insula | Ctx_AVI_L | 58 | -32 | 20 | 2 | 4.55 |
| Ventrolateral prefrontal cortex | Ctx_a9_46v_R | 66 | 42 | 46 | 10 | 4.47 |
| Ventrolateral prefrontal cortex | Ctx_a9_46v_L | 35 | -36 | 44 | 6 | 4.40 |
| Ventromedial prefrontal cortex | Ctx_p32_R | 47 | 8 | 42 | 0 | 4.27 |
| Anterior insula | Ctx_AVI_R | 16 | 32 | 20 | 2 | 4.19 |
| Posterior cingulate cortex | Ctx_RSC_R | 7 | 6 | -16 | 32 | 4.06 |
| Ventromedial prefrontal cortex | Ctx_10r_L | 13 | -12 | 38 | -8 | 4.05 |
| Parahippocampal cortex | Ctx_PeEc_R | 5 | 26 | -22 | -30 | 4.04 |
| Cerebellum | Cblm_VI_R | 11 | 34 | -50 | -28 | 4.02 |
| Thalamus/Pulvinar | Thal_Pulv | 10 | -12 | -32 | 0 | 4.00 |
| Medial temporal cortex | Ctx_PreS_R | 11 | 14 | -36 | 0 | 3.99 |
| Cerebellum | Cblm_CrusI_R | 10 | 40 | -40 | -38 | 3.99 |
| Orbitofrontal cortex | Ctx_OFC_L | 3 | -8 | 50 | -24 | 3.98 |
| Dorsolateral prefrontal cortex | Ctx_p9_46v_R | 9 | 46 | 34 | 24 | 3.72 |
| Precuneus | Ctx_POS2_R | 8 | 14 | -60 | 24 | 3.70 |
| Ventrolateral thalamus | Thal_VL | 7 | -12 | -12 | 14 | 3.65 |
| Cerebellum | Cblm_VI_L | 6 | -34 | -56 | -26 | 3.61 |
| Precuneus | Ctx_POS2_L | 3 | -12 | -68 | 36 | 3.55 |

***Negative weights***

| Name | Atlas label | Volume (voxels) | X | Y | Z | max(z) |
|---|---|---|---|---|---|---|

| | | | | | | |
|---|---|---|---|---|---|---|
| Visual cortex | Ctx_V2_L | 142 | -8 | -80 | -10 | -5.51 |
| Visual cortex | Ctx_V4t_L | 70 | -42 | -82 | 0 | -4.99 |
| Cerebellum | Cblm_CrusI_L | 148 | -28 | -80 | -30 | -4.93 |
| Cerebellum | Cblm_CrusI_R | 64 | 24 | -84 | -30 | -4.85 |
| Superior frontal cortex | Ctx_SFL_R | 47 | 4 | 6 | 68 | -4.85 |
| Paracentral lobule | Ctx_4_R | 37 | 6 | -30 | 60 | -4.83 |
| Superior parietal lobule | Ctx_7AL_L | 158 | -20 | -40 | 64 | -4.63 |
| Parahippocampal cortex | Ctx_VMV1_R | 59 | 20 | -44 | -10 | -4.60 |
| Visual cortex | Ctx_V3A_L | 94 | -12 | -86 | 24 | -4.51 |
| Superior temporal sulcus | Ctx_STSdp_L | 78 | -50 | -32 | -6 | -4.49 |
| Cerebellum | Cblm_V_L | 16 | -22 | -34 | -30 | -4.49 |
| Midcingulate cortex | Ctx_24dv_L | 21 | -8 | -2 | 40 | -4.40 |
| Parahippocampal cortex | Ctx_VMV1_L | 32 | -18 | -60 | -8 | -4.36 |
| Superior frontal cortex | Ctx_8BL_R | 35 | 2 | 50 | 46 | -4.28 |
| Visual cortex | Ctx_V4t_R | 48 | 44 | -78 | -2 | -4.23 |
| Visual cortex | Ctx_V3_R | 41 | 16 | -72 | -8 | -4.22 |
| Superior frontal cortex | Ctx_8BL_R | 15 | 4 | 30 | 62 | -4.20 |
| Superior parietal lobule | Ctx_7AL_R | 14 | 18 | -44 | 70 | -4.13 |
| Putamen | Putamen_Pp_R | 3 | 28 | -12 | 10 | -4.10 |
| Putamen | Putamen_Pp_L | 10 | -30 | -22 | 10 | -4.06 |
| Superior frontal cortex | Ctx_9a_L | 24 | -10 | 62 | 26 | -4.05 |
| Paracentral lobule | Ctx_SCEF_L | 11 | -8 | -6 | 66 | -4.05 |
| Visual cortex | Ctx_V3A_R | 31 | 20 | -90 | 18 | -4.00 |
| Frontal pole | Ctx_10d_R | 7 | 12 | 68 | 16 | -3.90 |
| Temporal pole | Ctx_PeEc_L | 4 | -28 | -4 | -34 | -3.87 |
| Visual cortex | Ctx_MT_R | 9 | 48 | -68 | 4 | -3.83 |
| Superior temporal gyrus | Ctx_PSL_L | 17 | -56 | -36 | 16 | -3.80 |
| Precuneus | Ctx_5L_R | 6 | 12 | -48 | 66 | -3.80 |
| Visual cortex | Ctx_V4_R | 8 | 32 | -78 | -10 | -3.79 |
| Superior temporal gyrus | Ctx_A4_L | 4 | -60 | -30 | 10 | -3.70 |
| Temporal pole | Ctx_TE1a_L | 4 | -52 | -6 | -32 | -3.70 |
| Superior temporal sulcus | Ctx_STV_R | 6 | 50 | -42 | 10 | -3.68 |
| Superior temporal gyrus | Ctx_PSL_R | 5 | 54 | -32 | 22 | -3.56 |

**Suppl. Table S5.** Significant positive and negative weights contributing to the k-marker for the parametric modulation by DELAY (FDR corrected q < 0.05 across the whole feature space, i.e. three concatenated gray-matter-masked whole brain maps, at least three contingent voxels).

***Positive weights***

| Name | Atlas label | Volume (voxels) | MNI Coordinates | | | max(z) |
|---|---|---|---|---|---|---|
| | | | X | Y | Z | |
| Visual cortex | Ctx_V3A_R | 131 | 12 | -86 | 20 | 5.30 |
| Ventromedial prefrontal cortex | Ctx_10v_R | 160 | 2 | 42 | -20 | 5.14 |
| Precuneus | Ctx_PCV_L | 102 | -10 | -54 | 46 | 4.75 |
| Superior temporal sulcus | Ctx_STSdp_L | 63 | -54 | -34 | 4 | 4.71 |
| Superior temporal gyrus | Ctx_TA2_R | 59 | 52 | -8 | -2 | 4.66 |
| Precentral gyrus | Ctx_3b_L | 214 | -34 | -26 | 62 | 4.62 |
| Angular gyrus | Ctx_PGi_R | 27 | 40 | -60 | 22 | 4.61 |
| Ventromedial prefrontal cortex | Ctx_9m_R | 37 | 16 | 48 | 6 | 4.58 |
| Putamen | Putamen_Pp_L | 35 | -32 | -14 | 2 | 4.43 |
| Caudate | Cau_L | 36 | -22 | 22 | 2 | 4.35 |
| Superior temporal gyrus | Ctx_TPOJ1_R | 29 | 50 | -30 | 8 | 4.34 |
| Angular gyrus | Ctx_PGi_L | 43 | -48 | -54 | 22 | 4.34 |
| Precentral gyrus | Ctx_4_L | 5 | -32 | -16 | 44 | 4.33 |
| Cingulate sulcus | Ctx_24dd_L | 49 | -6 | -10 | 48 | 4.29 |
| Subgenual cingulate cortex | Ctx_s32_L | 17 | -10 | 26 | -12 | 4.24 |
| Paracentral lobule | Ctx_5L_R | 28 | 10 | -40 | 62 | 4.16 |
| Midcingulate cortex | Ctx_24dv_R | 9 | 10 | -4 | 44 | 4.15 |
| Cerebellum | Cblm_CrusII_L | 3 | -12 | -84 | -44 | 4.04 |
| Caudate | Cau_R | 7 | 6 | 26 | -4 | 3.99 |
| Superior frontal gyrus | Ctx_9m_R | 21 | 4 | 56 | 38 | 3.97 |
| Midtemporal gyrus | Ctx_MT_R | 9 | 46 | -68 | 12 | 3.96 |
| Visual cortex | Ctx_V1_L | 14 | -14 | -82 | -4 | 3.94 |
| Cerebellum | Cblm_CrusII_R | 7 | 28 | -84 | -44 | 3.93 |
| Posterior insula/temporal operculum | Ctx_52_R | 16 | 40 | -22 | -2 | 3.91 |
| Superior temporal sulcus | Ctx_A5_L | 8 | -58 | 2 | -10 | 3.90 |
| Paracentral lobule | Ctx_5mv_R | 4 | 18 | -32 | 44 | 3.83 |
| Cerebellum | Cblm_CrusI_R | 10 | 40 | -74 | -34 | 3.82 |
| Midtemporal gyrus | Ctx_A5_L | 4 | -62 | -12 | -10 | 3.80 |
| Superior parietal lobule | No_label | 4 | -22 | -44 | 50 | 3.78 |
| Paracentral lobule | Ctx_5m_L | 11 | -10 | -44 | 64 | 3.77 |
| Paracentral lobule | Ctx_5mv_R | 7 | 16 | -24 | 44 | 3.72 |
| Inferior frontal cortex | Ctx_45_L | 5 | -54 | 26 | 14 | 3.72 |
| Precentral gyrus | Ctx_6d_L | 3 | -26 | -18 | 62 | 3.72 |
| Superior parietal lobule | Ctx_7Am_L | 5 | -4 | -66 | 62 | 3.70 |
| Visual cortex | Ctx_V2_L | 8 | -6 | -86 | 18 | 3.69 |
| Visual cortex | Ctx_IP0_R | 6 | 30 | -70 | 22 | 3.68 |
| Caudate | Cau_L | 4 | -18 | 8 | 18 | 3.68 |
| Posterior cingulate cortex | Ctx_24dd_R | 7 | 10 | -20 | 44 | 3.66 |

| | | | | | | |
|---|---|---|---|---|---|---|
| Superior parietal lobule | Ctx_2_L | 3 | -18 | -38 | 60 | 3.63 |
| Superior temporal sulcus | Ctx_STV_R | 3 | 58 | -34 | 8 | 3.57 |

***Negative weights***

| **Name** | **Atlas label** | **Volume (voxels)** | **X** | **Y** | **Z** | **max(z)** |
|---|---|---|---|---|---|---|
| Dorsolateral prefrontal cortex | Ctx_a9_46v_L | 76 | -40 | 50 | 14 | -5.20 |
| Brainstem | Bstem_Ponscd | 54 | 2 | -28 | -36 | -4.78 |
| Intraparietal sulcus | Ctx_7PC_L | 34 | -40 | -46 | 48 | -4.70 |
| Dorsolateral prefrontal cortex | Ctx_8C_R | 26 | 40 | 12 | 30 | -4.57 |
| Brainstem | Bstem_Ponscd_L | 69 | -12 | -38 | -36 | -4.54 |
| Paracentral lobule | Ctx_4_R | 12 | 6 | -30 | 78 | -4.41 |
| Inferior parietal lobule | Ctx_PFm_R | 23 | 54 | -44 | 52 | -4.39 |
| Cerebellum | Cblm_Dentate_R | 39 | 18 | -42 | -36 | -4.25 |
| Cerebellum | Cblm_IX_L | 51 | -6 | -54 | -36 | -4.23 |
| Medial frontal gyrus | Ctx_8BM_R | 45 | 0 | 22 | 42 | -4.17 |
| Orbitofrontal cortex | Ctx_a10p_R | 5 | 22 | 58 | -10 | -4.16 |
| Visual cortex | Ctx_V4_R | 19 | 30 | -86 | -10 | -4.11 |
| Paracentral lobule | Ctx_SFL_R | 10 | 4 | -2 | 76 | -4.10 |
| Thalamus | Thal_Pulv | 13 | -4 | -30 | 4 | -4.08 |
| Orbitofrontal cortex | Ctx_11l_R | 17 | 22 | 32 | -18 | -4.07 |
| Visual cortex | Ctx_LO2_L | 3 | -46 | -82 | -6 | -4.07 |
| Anterior insula/frontal operculum | Ctx_AVI_R | 3 | 36 | 26 | -4 | -4.06 |
| Dorsolateral prefrontal cortex | Ctx_p9_46v_R | 52 | 44 | 36 | 16 | -3.99 |
| Cerebellum | Cblm_CrusII_R | 3 | 8 | -86 | -28 | -3.92 |
| Frontal pole | Ctx_a10p_R | 9 | 28 | 66 | -6 | -3.85 |
| Thalamus | Thal_Pulv | 19 | 6 | -26 | 8 | -3.85 |
| Orbitofrontal cortex | Ctx_pOFC_R | 8 | 16 | 10 | -16 | -3.84 |
| Midcingulate cortex | Ctx_a32pr_L | 4 | -8 | 34 | 24 | -3.84 |
| Visual cortex | Ctx_V4_L | 10 | -34 | -90 | -6 | -3.80 |
| Frontal pole | Ctx_a10p_L | 3 | -20 | 60 | -10 | -3.80 |
| Visual cortex | Ctx_V3CD_L | 9 | -44 | -86 | 12 | -3.79 |
| Superior parietal lobule | No_label | 3 | 28 | -38 | 76 | -3.79 |
| Posterior cingulate cortex | Ctx_31a_L | 9 | 0 | -32 | 44 | -3.74 |
| Inferior parietal lobule | Ctx_PFm_L | 4 | -50 | -48 | 52 | -3.69 |
| Frontal pole | Ctx_p10p_L | 3 | -30 | 66 | 0 | -3.65 |
| Visual cortex | Ctx_VVC_R | 4 | 30 | -62 | -14 | -3.63 |
| Nucleus Accumbens | NAC_L | 6 | -10 | 4 | -18 | -3.57 |
| Nucleus Accumbens | NAC_L | 3 | -8 | 12 | -10 | -3.52 |

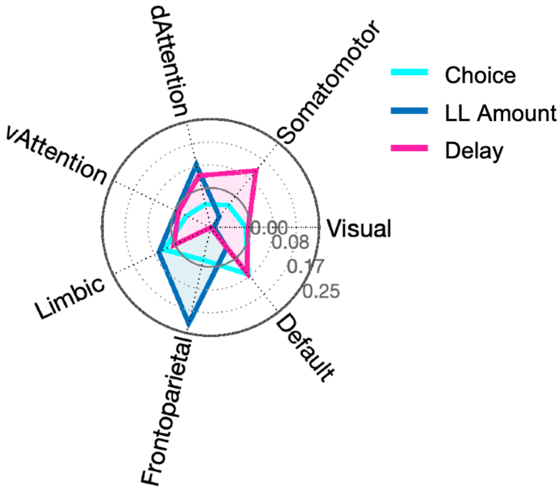


**Fig. S6**. **Spatial similarity of the *k-marker* with seven established large-scale functional resting-state networks (Yeo et al., 2011).** The spatial correlation (Pearson's *r*) of the *k*-marker patterns with each network (Yeo et al., 2011) with each component map of the *k*-marker (Choice, LL Amount, Delay) is displayed in a radar plot. The *k*-marker pattern for the Choice contrast did not correlate strongly with any given resting state network (highest of $r = 0.074$ with the Default network). The pattern for the LL-Amount contrast correlated positively with the Frontoparietal network (FPN) (r = 0.22), indicating that higher activity in this network—often associated with cognitive control—for greater LL amounts predicts *greater* delay discounting. Negative correlations were seen with the Visual and Somatosensory networks, in line with the findings reported above. In contrast, the pattern for the Delay contrast correlated positively with the Somatomotor network (r = 0.12) and negatively with the FPN (r = -0.15). This suggests that networks may contribute with different signs to different aspects of decision-making. While increased FPN activity for increasing LL amounts predicts greater discounting, increased FPN activity for increasing delays predicts *lower* discounting. Paralleling the meta-analytical similarity results in the main text, this suggests that the relationship between control-related activity and discounting is less clear-cut than previously thought.
